# Derivation of Poly-Methylomic Profile Scores for Schizophrenia

**DOI:** 10.1101/607309

**Authors:** Oliver J. Watkeys, Sarah Cohen-Woods, Yann Quidé, Murray J. Cairns, Bronwyn Overs, Janice M. Fullerton, Melissa J. Green

## Abstract

Schizophrenia (SZ) and bipolar disorder (BD) share numerous clinical and biological features as well as environmental risk factors that may be associated with altered DNA methylation. In this study we sought to construct a Poly-Methylomic Profile Score (PMPS) for SZ, representing the degree of epigenome-wide methylation according to previously published findings; we then examined its association with SZ and BD in an independent sample. DNA methylation for 57 SZ, 59 BD cases and 55 healthy controls (HCs) was quantified using the Illumina 450K methylation beadchip. We constructed five PMPSs for different p-value thresholds using summary statistics reported in a large epigenome-wide schizophrenia case-control association study, weighted by individual CpG effect sizes. All SZ PMPSs were significantly elevated in SZ cases relative to HCs, with the score calculated at the most stringent threshold accounting for the greatest amount of variance in SZ (compared to other PMPSs derived at more inclusive *p*-value thresholds). However, none of the PMPSs were associated with BD, or a combined cohort of BD and SZ cases relative to HCs. Results demonstrating elevated PMPSs in SZ relative to BD did not survive correction for multiple testing. PMPSs were also not associated with positive or negative symptom severity. That this SZ-derived PMPSs was elevated among SZ, but not BD participants, suggests that epigenome-wide methylation patterns associated with schizophrenia may represent distinct pathophysiology that is yet to be elucidated. Whether this PMPS may be associated with neuroanatomical or other biological endophenotypes relevant to SZ and/or BD remains to be determined.

## 1. Introduction

Owing to recent advances in genomic technologies and collaboration among global consortia, schizophrenia (SZ) and bipolar disorder (BD) have been shown to share a significant proportion of genetic liability (Hamshere et al., 2011; Ivleva et al., 2010; Purcell et al., 2009). For example, the degree to which multiple sites of genetic variation contribute to risk for SZ and BD can be summarised using ‘polygenic risk scores’, calculated as the sum of alleles associated with a particular trait, weighted by their respective effect sizes (Purcell et al., 2009). While the majority of these genetic variants would individually fail to reach even nominal levels of statistical significance, collectively they may account for a significant proportion of the variance in manifestation of SZ and BD (Purcell et al., 2009). However, there remains a significant amount of unexplained variance in shared genetic and environmental risk for these disorders (Lichtenstein et al., 2009), some of which may be accounted for by epigenetic processes associated with shared environmental risk factors such as childhood trauma, cannabis use, obstetric complications (Bourque et al., 2011; Buoli et al., 2016; Cannon et al., 2002; Kraan et al., 2016; Laursen et al., 2007).

Epigenetic processes, such as DNA methylation, refer to functionally significant alterations in the genome that do not alter the nucleotide sequence (Hannon et al., 2016; Montano et al., 2016; Pries et al., 2017). DNA methylation involves the addition of a methyl-group to the carbon-5 position of a cytosine residue (Razin and Cedar, 1991): this may be associated with increased or decreased gene expression depending on the genomic region in which it occurs (Jones, 2012; Watkeys et al., 2018). Both candidate- and genome-wide studies have revealed differential methylation of various sites in SZ (Hannon et al., 2016; Montano et al., 2016; Ruzicka et al., 2015; Yoshino et al., 2016), with some studies suggesting overlapping patterns of DNA methylation in BD (Dempster et al., 2011; Sugawara et al., 2018), although there have been few replications at individual CpG sites (Pries et al., 2017; Teroganova et al., 2016).

A recently published epigenome-wide association study (EWAS) (Hannon et al., 2016) of blood-derived DNA methylation patterns reported 14 differentially methylated CpG sites that were robustly associated with SZ in both a large discovery sample (comprising 353 SZ cases and 322 healthy controls) and a replication sample (comprising 414 SZ cases and 433 healthy controls), at a stringent significance level (*p* < 1 × 10^−7^). A larger number of CpGs (n=213) were associated with SZ in both the discovery and replication case-control samples at a more liberal significance threshold (*p* < 1×10^−5^). Hannon et al.’s gene ontology analysis suggested that genes containing CpG sites associated with SZ were implicated in inflammatory processes and brain development. Importantly, Hannon et al. (Hannon et al., 2016) reported effect sizes and *p*-values for all CpG sites differentially methylated in SZ for all samples (with replication *p*-values ranging from 4.42×10^−18^ to 0.991), and included covariates (age, sex, technical batch effects, as well as methylation-derived estimates of cell composition and smoking status). A comparably sized study published in the same year reported only CpG sites significant at an FDR corrected threshold of *p*<0.2 in their replication sample (number CpGs = 172) (Montano et al., 2016).

In this study, we adapted methodology used to calculate PRS from GWAS summary statistics, to estimate a series of poly-methylomic profile scores (PMPSs) for SZ. The calculation of these scores was based on discovery effect sizes of individual differentially methylated CpG sites at various *p*-value thresholds as reported by Hannon et al. (2016). We then established the extent to which these PMPSs were associated with an independent sample of cases with SZ, BD, and the combined SZ-BD group (each compared to healthy controls; HC), and directly compared PMPSs between SZ and BD. We also explored associations with positive and negative symptoms of psychosis for the PMPS that was most strongly associated with SZ.

## 2. Experimental/Materials and methods

Study procedures were approved by the UNSW Human Research Ethics committees (HC12384), St. Vincent’s Hospital (HREC/10/SVH/9), and the South East Sydney and Illawarra Area Health Service (HREC 09/081).

### 2.1. Participants

Peripheral blood samples were obtained from a total of 191 study participants (64 SZ, 64 BD, 60 HC). Following the application of exclusion criteria and QC processes (see below) the final sample (N=171) comprised 57 people who met ICD-10 (World Health Organisation, 2004) criteria for SZ or schizoaffective disorder (referred to as SZ), 59 individuals with BD-I disorder (BD), and 55 HCs. Clinical participants were recruited from outpatient services of the South Eastern Sydney-Illawarra Area Health Service (SESIAHS), the Australian Schizophrenia Research Bank (Loughland et al., 2010), and the Sydney BD Clinic (Mitchell et al., 2009). Healthy controls were recruited from the local community. Study exclusion criteria included current neurological disorder, current substance abuse or dependence, and/or electroconvulsive treatment in the past six months. Healthy controls were also required to have no lifetime history of a DSM-IV Axis-I diagnosis (American Psychiatric Association, 1994) confirmed using the Mini-International Neuropsychiatric Interview (Sheehan et al., 1998), and no history of psychotic disorders among first-degree biological relatives.

### 2.2. Cognitive and symptom measures

Estimates of premorbid and current intelligence quotients (IQ) were calculated using the Wechsler Test of Adult Reading (Wechsler, 2001), and the Wechsler Abbreviated Scale of Intelligence (Wechsler, 1999), respectively. The Scales for the Assessment of Positive Symptoms (SAPS) and Negative Symptoms (SANS) were used to assess symptom severity (Andreasen, 1984a, b).

### 2.3. Methylation quantification and quality control

Methylation quantification was undertaken on DNA samples from 191 participants at the Institute for Molecular Bioscience, University of Queensland (UQ), according to standard manufacturer’s instructions using the Illumina 450K BeadChip. Samples were randomised on the plate to avoid bias due to chip or position on chip. Quantification was completed using the Qubit fluorimeter, and bisulfite conversions were completed using the EZ-96 DNA Methylation kit according to the manufacturers protocol (Zymo Research, Orange, CA, USA), and assessed on the Epoch Microplate Spectrophotometer (Biotek, Winooski, VT). All chips were imaged and analysed for DNA methylation status on the Illumina iScan system using the GenomeStudio methylation module software (Illumina, San Diego, CA, USA). This produced a methylation score as a continuous β-value representing the ratio of methylated (M) probe intensity relative to total intensity of methylated and unmethylated (U) probes defined as β=*M*/(*M*+*U*). Raw and total probe intensities were extracted.

Of the 191 samples, 94 had DNA derived from whole blood, 94 firstly underwent ficoll density gradient separation with DNA extracted from the cells in the ficoll/PBS/plasma layer, and 3 were derived from the isolated lymphoblast cell population after ficoll separation; each with different cellular compositions in each tissue source. Ten samples were excluded from methylation QC analyses owing to their lymphoblast origin (3 samples) or having undergone 450k hybridisation in a different batch (7 samples); 181 DNA samples (whole blood or ficoll-treated) were included in methylation QC procedures.

The R package ‘meffil’ (Min et al., 2017) was used to extract signal intensities and perform background correction and normalisation of methylation beta values for individual CpG sites. Our QC procedure involved excluding individual subjects for whom: (i) more than 10% of probes had less than three beads (0 samples), (ii) the predicted median methylation signal was in excess of three standard deviations from the regression line generated from methylated vs. unmethylated signal intensities (1 sample), (iii) a gender mismatch (0 samples) or XY outlier status was detected (2 samples), (iv) methylation signal intensities deviated excessively from mean values for control probes (4 samples), (v) more than 10% of probes had a detection *p*-value > 0.01 (0 samples), or (vi) a genotype mismatch with genotypes derived from Illumina psych-chip was detected (1 sample). Individual probes were excluded if (i) >10% of samples showed detection *p*-values > 0.01 for that probe (n = 137), (ii) >10% of samples demonstrated less than 3 beads at a given probe (n = 141), or (iii) the probe was previously demonstrated to be cross-reactive (n = 29 233) (Chen et al., 2013). After application of these QC procedures, 456 041 CpG sites remained for analysis in the 171 participants. However, in the current study we only included the 1051 independent CpG sites demonstrating a replication *p*-value < 0.5 in Hannon et al.’s (Hannon et al., 2016) first replication cohort.

Blood cell count estimates were derived from methylation data for Bcells, CD4T, CD8T, granulocytes, monocytes, and natural killer cells using meffil’s gse35069 profile references. Shinymethyl (Fortin et al., 2014) was used to conduct principle components analysis with the final QCed data. We found two components significantly associated with ethnicity (*p* < 0.05), which were included as covariates in all subsequent analysis.

### 2.4. Poly-Methylomic Profile Score (PMPS)

PMPSs were calculated utilising data from multiple CpG sites previously reported as demonstrating an association with SZ (Hannon et al., 2016), weighted on the basis of discovery sample effect sizes. Sites were included in the PMPSs if they replicated across the two cohorts tested by Hannon et al. (2016) at five different *p*-value thresholds (1 × 10^−7^, 1 × 10^−5^, 0.001, 0.05, 0.5). The use of multiple *p*-value thresholds is standard in the construction of polygenic risk scores, in order to identify the threshold that enables optimal prediction (Purcell et al., 2009). The five *p*-value thresholds included 14, 210, 359, 684, and 1051 CpG sites in each PMPS, respectively. The first two scores were calculated based on CpG sites which replicated at their respective *p*-value threshold across both the discovery and first replication cohort. In contrast, the final three PMPSs were calculated based on sites that evidenced at least a discovery threshold (*p* < 1 × 10^−5^) association with schizophrenia and replicated at each of the three thresholds listed above, respectively (Hannon et al., 2016). Each PMPS score was calculated using previously described methods (Shah et al., 2015) as the sum of weighted beta values associated with SZ, or:

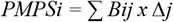

Where PMPSi is the polymethylomic profile score for the i^th^ participant, Bij is the beta value for the i^th^ participant at the j^th^ CpG site, and Δj is the effect size for the j^th^ CpG site.

To ensure that each PMPS was not inflated by correlated methylation among CpGs, the independence of CpG sites meeting thresholds for inclusion in any PMPS was assessed via correlation analyses. In the event that two or more CpG sites within 500 base-pairs (bp) of each other demonstrated a Pearson’s correlation coefficient > 0.1, a single site demonstrating the strongest association with SZ (as indicated by replication *p*-values from Hannon et al. (2016) was retained for PMPS calculation, whilst the other correlated sites (n=10) were excluded (see Supplementary Table 1) (Huynh et al., 2013; Ong and Holbrook, 2013; Shah et al., 2015).

As smoking behaviour influences methylation (Zeilinger et al., 2013), we used the method described above to estimate a methylation smoking score, using smoking-related methylation effect sizes for CpG sites demonstrating a significant replicated association with smoking status (*p*-value < 5 × 10^−5^) reported previously (Zeilinger et al., 2013). We validated this score using smoking data collected for our clinical participants; the smoking score was significantly associated with both lifetime smoking (*B* = 0.880, *OR* = 2.410, *p* = 6.49 × 10^−5^) and Fagerstrom test for nicotine dependency (Heatherton et al., 1991) (*B* = 0.395, *OR* = 1.484, *p* = 0.004).

### 2.5. Statistical Analyses

An initial exploratory analysis investigated whether patterns of methylation for individual CpG sites reported by Hannon et al. (2016) were differentially methylated in the current SZ sample. Beta values for each CpG site were used as dependent variables and clinical status (SZ/HC) as the main explanatory variable. For this and all subsequent analyses, covariates of age, sex, tissue source (ficoll or whole blood), estimated methylation-derived cell counts, two methylation-derived ethnicity principle components, and an estimated methylation smoking score were included as fixed factors, and 450k slide ID as a random factor. Individual CpG sites were only considered to be significantly associated with SZ if they evidenced a False Discovery Rate (FDR) corrected *q*-value < 0.05.

A series of logistic regression analyses were then performed to assess if the PMPSs were associated with: (i) SZ (relative to HC); (ii) BD (relative to HC); (iii) the combined BD and SZ group (relative to HC); and (iv) SZ (relative to BD). The PMPSs were converted to z-scores before analysis so that regression coefficients and effect sizes would be comparable between models. FDR correction was applied across the five PMPSs assessed within each group comparison. In addition, two linear regression analyses were used to assess if the optimal PMPS was associated with SAPS and SANS global subscale scores respectively.

To investigate any potential confounding effects of medication, we ran three linear regression models (one model each for BD, SZ, and the combined SZ-BD cohort), with the PMPS that explained the most variance in SZ as the dependent variable, and imipramine (antidepressant) equivalency dosage, chlorpromazine (antipsychotic) equivalency dosage, and mood stabiliser usage included as predictors in addition to covariates used in all other analyses.

## 3. Results

Demographic information for included participants is summarised in Table 1.

**Table 1.** Sample demographics. Legend: Demographics statistics for each group (schizophrenia, bipolar disorder, and healthy controls)

| Variable | HC | BD | SZ | Group difference statistics |
| --- | --- | --- | --- | --- |
| N (% Male) | 55 (49.09) | 59 (28.81) | 57 (61.40) | $\chi^2 = 12.661, p = 0.002$ |
| Mean Age in years (SD) | 36.32 (11.78) | 37.66 (12.56) | 43.02 (11.12) | $F(2,168) = 5.061, p = 0.007, \eta_p^2 = 0.057$ |
| Mean Education in years (SD) | 16.84 (2.46) | 15.68 (2.79) | 14.23 (2.49) | $F(2,168) = 14.289, p = 1.858 \times 10^{-6}, \eta_p^2 = 0.145$ |
| Mean WTAR (SD) | 110.62 (10.30) | 105.02 (14.78) | 100.21 (14.43) | $F(2,167) = 8.495, p = 3.066 \times 10^{-4}, \eta_p^2 = 0.092$ |
| Mean WASI (SD) | 117.74 (12) | 114.82 (11.39) | 107.7 (14.077) | $F(2,124) = 7.121, p = 0.001, \eta_p^2 = 0.103$ |
| Mean SAPS (SD) | NA | 6.75 (8.19) | 17.75 (14.61) | $F(1,111) = 24.469, p = 2.706 \times 10^{-6}, \eta_p^2 = 0.181$ |
| Mean SANS (SD) | NA | 15.3 (15.83) | 29.84 (17.26) | $F(1,110) = 21.605, p = 9.361 \times 10^{-6}, \eta_p^2 = 0.164$ |
| Mean Age of illness onset | NA | 23.42 (9.70) | 25.04 (8.51) | $F(1,105) = 0.831, p = 0.364, \eta_p^2 = 0.008$ |
| Mood stabiliser usage (%; BD vs. SZ) | NA | 45 (76.27) | 12 (21.05) | $\chi^2 = 33.195, p = 8.34 \times 10^{-9}$ |
| Mean Imiprimine-equivalent dosage (SD) | NA | 26.06 (60.05) | 63.69 (121.99) | $F(1,114) = 4.491, p = 0.036, \eta_p^2 = 0.038$ |
| Mean Chlorpromazine-equivalent dosage (SD) | NA | 194 (571.09) | 476.9 (886.43) | $F(1,114) = 4.204, p = 0.043, \eta_p^2 = 0.036$ |

HC= healthy controls; BD= bipolar disorder; SZ= schizophrenia; WTAR=Wechsler Test of Adult Reading; WASI=Wechsler Abbreviated Scale of Intelligence; SAPS= Scale for Assessment of Positive Symptoms; SANS=Scale for Assessment of Negative Symptoms; SD=standard deviation

### 3.1. Differential methylation analysis

Of 1051 CpG sites included for analysis in the present study, only 79 were associated with SZ in our sample at a nominal *p*-value threshold (*p* < 0.05), with none surviving FDR correction for multiple testing (FDR < 0.05; see Supplementary Tables 2 and 3 for full results). Of the 14 CpG sites most robustly associated with SZ in both Hannon et al.’s (2016) discovery and replication cohorts (*p* < 1×10^−7^), only two sites (cg13803727 and cg27541604) demonstrated nominally significant associations with SZ in our cohort (*p* < 0.05) (see Supplementary Table 2). There was 100% consistency at these 14 sites between the direction of hypo-/hyper-methylation reported in Hannon et al.’s (Hannon et al., 2016) discovery cohort and our own sample. The consistency in the direction of methylation between Hannon et al. (2016) discovery cohort and our own (81.42%, 74.93%, 64.91%, and 58.23%) tended to decrease at more liberal p-value thresholds (1×10^−5^, 0.001, 0.05, and 0.5, respectively).

### 3.2. PMPS model comparisons in association with SZ

All five PMPSs were significantly associated with SZ relative to HC both before and after FDR correction (see Table 2). All models explained a similar amount of the variance in SZ (58-60%), however models incorporating PMPSs based on more liberal *p*-value inclusion thresholds tended to explain less of the variance (see Table 2). The PMPS that explained the greatest amount of variance in SZ diagnosis incorporated the 14 CpG sites which were those most robustly associated with SZ across both discovery and replication samples from Hannon et al. (2016) (Nagelkerke Pseudo R^2^ = 0.595). Within this model, SZ cases showed heightened PMPS relative to controls (*B* = 1.263, *OR* = 3.538, 95% CI=1.227-10.197, *p* = 0.018; see Figure 1).

**Figure 1.**
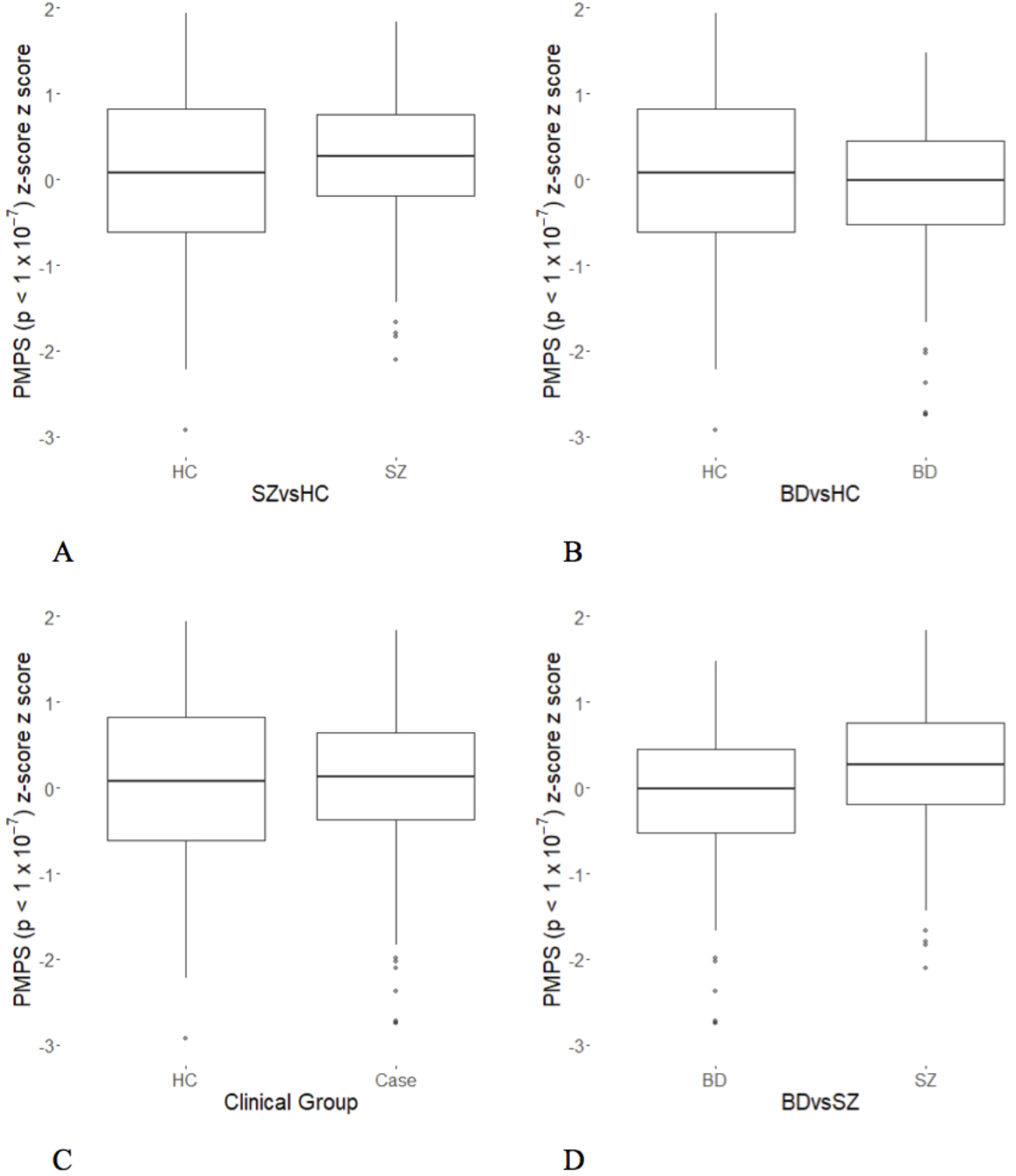
Group comparisons on raw PMPS score. Legend: Boxplots showing group differences on the PMPS calculated at the most stringent *p*-value threshold (p 1 × 10^−7^) between: A) schizophrenia cases and healthy controls, B) bipolar disorder cases and healthy controls, C) combined clinical cases (SZ+BD) and healthy controls, and D) bipolar disorder and schizophrenia.

**Table 2.** Associations between each of the five PMPS scores and clinical groups. Legend: Results of logistic regression analyses comparing various groups on each PMPS.

| Group comparison | PMPS p-value threshold | # *CpGs* | *B* | *OR* | CI 95% | *p* | FDR q value | Pseudo R<sup>2</sup> |
| --- | --- | --- | --- | --- | --- | --- | --- | --- |
| SZ vs. HC | 1e-07 (discovery and replication) | 14 | 1.263 | 3.538 | 1.227-10.197 | 0.018 | 0.029 | 0.5958 |
|  | 1e-05 (discovery and replication) | 210 | 1.127 | 3.086 | 1.166-8.166 | 0.021 | 0.029 | 0.5934 |
|  | 0.001 (replication) | 359 | 0.975 | 2.651 | 1.091-6.441 | 0.029 | 0.029 | 0.5886 |
|  | 0.05 (replication) | 684 | 0.923 | 2.516 | 1.126-5.622 | 0.022 | 0.029 | 0.5919 |
|  | 0.5 (replication) | 1051 | 0.803 | 2.233 | 1.080-4.617 | 0.028 | 0.029 | 0.5883 |
| BD vs. HC | 1e-07 (discovery and replication) | 14 | -0.077 | 0.926 | 0.423-2.024 | 0.845 | 0.979 | 0.3998 |
|  | 1e-05 (discovery and replication) | 210 | -0.010 | 0.990 | 0.472-2.078 | 0.979 | 0.979 | 0.3995 |
|  | 0.001 (replication) | 359 | -0.130 | 0.878 | 0.424-1.817 | 0.722 | 0.979 | 0.4006 |
|  | 0.05 (replication) | 684 | -0.273 | 0.761 | 0.379-1.530 | 0.437 | 0.979 | 0.4045 |
|  | 0.5 (replication) | 1051 | -0.281 | 0.755 | 0.381-1.495 | 0.414 | 0.979 | 0.4051 |
| Case vs. Control | 1e-07 (discovery and replication) | 14 | 0.472 | 1.603 | 0.875-2.935 | 0.123 | 0.238 | 0.3246 |
|  | 1e-05 (discovery and replication) | 210 | 0.465 | 1.592 | 0.911-2.783 | 0.100 | 0.238 | 0.3269 |
|  | 0.001 (replication) | 359 | 0.381 | 1.464 | 0.860-2.494 | 0.157 | 0.238 | 0.3223 |
|  | 0.05 (replication) | 684 | 0.304 | 1.355 | 0.827-2.222 | 0.224 | 0.238 | 0.3188 |
|  | 0.5 (replication) | 1051 | 0.284 | 1.328 | 0.826-2.137 | 0.238 | 0.238 | 0.3183 |
| SZ vs. BD | 1e-07 (discovery and replication) | 14 | 0.852 | 2.345 | 0.970-5.666 | 0.055 | 0.055 | 0.5179 |
|  | 1e-05 (discovery and replication) | 210 | 0.834 | 2.303 | 1.035-5.121 | 0.038 | 0.055 | 0.5232 |
|  | 0.001 (replication) | 359 | 0.766 | 2.151 | 1.021-4.532 | 0.041 | 0.055 | 0.5219 |
|  | 0.05 (replication) | 684 | 0.789 | 2.202 | 1.094-4.433 | 0.025 | 0.055 | 0.5292 |
|  | 0.5 (replication) | 1051 | 0.655 | 1.925 | 0.998-3.714 | 0.048 | 0.055 | 0.5198 |

### 3.3. SZ-PMPS in association with BD and the cross-disorder sample

In contrast to the SZ-HC PMPS models (described above), models comparing BD and the combined “psychosis spectrum” group (SZ + BD = 116 cases) to HCs, explained considerably less of the variance (39-41% and 31-33%, respectively). Furthermore, none of the SZ PMPSs within these models were significantly associated with a diagnosis of BD (relative to HCs), nor were they significantly associated with the combined “psychosis spectrum” group relative to HCs either before or after FDR correction. Conversely, a considerable degree of the variance in BD versus SZ status was accounted for by the PMPS models (51-53%). While there was no significant difference between BD and SZ participants on the PMPS calculated at the most stringent *p*-value threshold (*p* < 1 × 10^−7^) (*B* = 0.852, *OR* = 2.345, 95% CI = 0.970-5.666, *p* = 0.055) all other PMPS scores were significantly elevated in SZ relative to BD. However, none of these associations withstood FDR correction (see Table 2).

### 3.4. Associations between optimal PMPS and positive and negative symptoms of psychosis

There were no relationships between the optimal PMPS and any subscales of the SAPS and SANS, respectively (see Table 3).

**Table 3.** PMPSs and SAPS/SANS subscale scores. Legend: Results for regression models examining association between PMPS (p < 1 × 10^−7^) and SANS/SAPS global subscales.

| **SANS/SAPS subscale score** | ***B*** | ***OR*** | ***95% CI*** | ***p*** |
| --- | --- | --- | --- | --- |
| SANS Affective Flattening | 0.117 | 1.124 | 0.933-1.354 | 0.216 |
| SANS Alogia | -0.047 | 0.954 | 0.797-1.142 | 0.607 |
| SANS Avolition | 0.056 | 1.057 | 0.900-1.242 | 0.491 |
| SANS Asociality | 0.029 | 1.030 | 0.880-1.204 | 0.712 |
| SANS Attention | -0.083 | 0.920 | 0.784-1.079 | 0.302 |
| SAPS Hallucinations | 0.033 | 1.034 | 0.874-1.222 | 0.696 |
| SAPS Delusions | 0.114 | 1.121 | 0.918-1.369 | 0.258 |
| SAPS Bizarre behaviour | 0.004 | 1.004 | 0.800-1.259 | 0.975 |
| SAPS Thought disorder | -0.055 | 0.946 | 0.827-1.082 | 0.414 |

### 3.5. Potential effects of medication on PMPS scores

Irrespective of clinical group (SZ, BD, or cross-diagnostic “psychosis spectrum” group), all linear regression analyses indicated that medication usage was not significantly associated with the optimal PMPS (all *p* > 0.05; see Table 4).

**Table 4.** PMPSs and Medication. Results from linear regression analyses investigating relationship between PMPS (1 × 10^−7^) for SZ and medication use in cases.

| Cohort and Medication | *B* | *OR* | *95% CI* | *p* |
| --- | --- | --- | --- | --- |
| **SZ** |  |  |  |  |
| Imipramine Equivalent Dosage | -0.001 | 0.999 | 0.996-1.002 | 0.378 |
| Chlorpromazine Equivalent Dosage | 0.000 | 1.000 | 1.000-1.001 | 0.115 |
| Mood Stabiliser Usage | 0.217 | 1.242 | 0.482-3.198 | 0.641 |
| **BD** |  |  |  |  |
| Imipramine Equivalent Dosage | 0.003 | 1.003 | 0.996-1.009 | 0.419 |
| Chlorpromazine Equivalent Dosage | 0.000 | 1.001 | 0.999-1.001 | 0.807 |
| Mood Stabiliser Usage | -0.055 | 0.946 | 0.485-1.848 | 0.867 |
| **Combined psychosis group (SZ+BD)** |  |  |  |  |
| Imipramine Equivalent Dosage | 0.001 | 1.001 | 0.999-1.002 | 0.460 |
| Chlorpromazine Equivalent Dosage | 0.000 | 1.000 | 1.000-1.000 | 0.332 |
| Mood Stabiliser Usage | -0.202 | 0.817 | 0.586-1.138 | 0.228 |

Note: The optimal PMPS score for SZ was based on the top 14 CpG sites associated with SZ as reported by Hannon et al.<sup>5</sup>

## 4. Discussion

Here we demonstrate the construction of a Poly-Methylomic Profile Score for schizophrenia based on differentially methylated CpG sites reported in a recent EWAS in schizophrenia (2016); we demonstrate significantly higher scores in an independent schizophrenia sample, relative to healthy controls, at both stringent and increasingly liberal thresholds of significance. Although we found little evidence for significant differences in methylation at those individual CpG sites in this small schizophrenia sample, likely due to limited power, there was 100% consistency in the direction of DNA hyper-/hypo-methylation in our schizophrenia sample among the 14 CpG sites previously reported as most strongly associated with schizophrenia (Hannon et al., 2016). Interestingly, schizophrenia PMPSs were not associated with BD or the combined psychosis group compared to healthy controls, suggesting that the SZ-derived PMPS may reflect distinct pathophysiology in SZ. Furthermore, whilst the PMPS differences between SZ and BD were not significant after FDR correction, these associations were only just above the threshold of statistical significance. Whilst it is possible that this indicates shared methylation patterns between SZ and BD, it is also possible that our study was underpowered to detect true differences in the methylation profiles between SZ and BD.

The capacity to demonstrate associations between the constructed poly-methylomic scores and schizophrenia is consistent with the idea that methylation changes in schizophrenia span multiple CpG sites of small effect throughout the genome, and that collectively, the summed effects across multiple CpG sites should explain a larger proportion of the variance in the manifestation of schizophrenia than individual CpG sites alone. Notably, the amount of variance explained by the PMPSs decreased as increasingly liberal *p*-value thresholds were adopted for score calculation. This stands in stark contrast to genetic polygenic risk scores, which tend to explain more of the variance when more liberal *p*-value thresholds are utilised (Purcell et al., 2009). This difference may reflect the smaller EWAS cohorts compared to those used to generate polygenic risk scores for schizophrenia (Purcell et al., 2009). However, additional consideration should also be given to the increased extent to which DNA methylation is susceptible to confounding factors. That is, whilst genetic sequence variants are inherited directly from parents and remain stable across the lifespan (except in the case of de novo events), epigenetic processes are susceptible to environmental modification and change over time (Dempster et al., 2013). These factors make it inappropriate to characterise a trait-associated methylation score as indexing *risk* for the trait in question. Rather, any such score might be instead regarded as summarising the methylation *profile* associated with a given trait. Indeed, such scores have already been calculated for phenotypes such as body-mass index (BMI) and environmental exposures such as smoking status, education, and alcohol consumption (Elliott et al., 2014; Hamilton et al., 2018; McCartney et al., 2018). Our study reflects the first attempt to derive such scores from methylome-wide CpG data for established schizophrenia. Application of this profile to first-episode psychosis samples, and the further derivation of PMPSs from EWASs conducted in first-episode psychosis and other medication naive high-risk samples, will be useful to understand its potential biological relevance to disease risk and the emerging pathophysiology of schizophrenia.

Interestingly, none of the SZ-derived PMPS scores were associated with BD or the cross-disorder psychosis group relative to HCs, suggesting that the methylation patterns summarised by these scores may reflect distinct schizophrenia pathophysiology. This was somewhat unexpected given increasing evidence of biological overlap between schizophrenia and BD (Dempster et al., 2011; Ivleva et al., 2010; Laursen et al., 2007; Purcell et al., 2009; Sugawara et al., 2018). Furthermore, whilst PMPSs were not significantly elevated in SZ compared to BD after correcting for multiple testing, this could reflect lack of power. Larger discovery EWASs may be required to identify CpGs that are differentially methylated in both conditions. It would also be interesting to determine if a PMPS calculated using discovery data from a BD cohort is associated with both BD and SZ, however large discovery EWAS datasets in bipolar disorder are currently lacking.

Several limitations should be considered when interpreting these results. First, the reference profile used for estimating cellular composition was based on a small sample of men (Reinius et al., 2012). Whilst this method is widely used, questions remain regarding the validity of this measure as a means of estimating cellular composition in the broader population. Other potential sources of variation in DNA methylation include participants’ smoking behaviours (Zeilinger et al., 2013) and body mass index (BMI) (McCartney et al., 2018). We partially addressed this by estimating a methylation-based smoking score for use as a covariate (Elliott et al., 2014), since smoking status can be accurately estimated using methylation profile scores (Elliott et al., 2014). However, a methylation-derived score of BMI was not estimated, as there is substantially less reliability where BMI estimation is concerned (McCartney et al., 2018), and phenotypic BMI data was not available for our sample. Regarding potential medication effects, we saw no significant associations between PMPS for schizophrenia and antipsychotic and antidepressant dosages, or mood stabiliser use, but caution that further research in medication-naïve cohorts is warranted. It is also important to note that the tissue specificity inherent to DNA methylation patterns (Davies et al., 2012; Murphy et al., 2005; van den Oord et al., 2016; Walton et al., 2016; Xin et al., 2010) which are highly influenced by cellular composition (Jaffe and Irizarry, 2014), limits the generalisability of methylation patterns obtained from blood. Replicating the methods described here using discovery/replication cohorts derived from brain tissue will be important to understand the significance of methylation patterns over the course of illness.

In conclusion, we have developed a novel method for calculating a poly-methylomic profile of schizophrenia based on published EWAS data; all profile scores derived at various *p*-value thresholds were associated with SZ, but not with BD, or the cross-disorder sample of SZ and BD cases, relative to healthy participants. The PMPS has potential utility for estimating how epigenetic patterns across multiple differentially methylated sites might be related to intermediate (endo)phenotypes of schizophrenia and related disorders, such as cognitive functioning and brain structural and functional abnormalities. Future studies in larger independent samples should be mindful of potential demographic and experimental confounders of DNA methylation, such as tissue source of DNA, cellular composition, smoking status, BMI, age and medication exposure. Finally, efforts should be made to assess the extent to which blood-based PMPSs for schizophrenia index methylation patterns in the brain, and the extent to which these markers can shed light on the pathophysiology of psychosis and related disorders.

