## Supplementary Tables 1 and 2 for "Derivation of Poly-Methylomic Profile Scores for Schizophrenia"

**SUPPLEMENTARY MATERIALS**

Watkeys, Oliver J. (BSc. Honours) ^a, b^

Cohen-Woods, Sarah (PhD) ^c, d, e^

Quidé, Yann (PhD) ^a, b^

Cairns, Murray J. (PhD) ^f^

Bronwyn Overs (BPsych. Honours) ^b^

Fullerton, Janice M. (PhD) ^b, g^

Green, Melissa J. (PhD)* ^a, b^

^a^ School of Psychiatry, University of New South Wales (UNSW Sydney), Sydney, NSW, Australia

^b^ Neuroscience Research Australia, Sydney, NSW, Australia

^c^ Discipline of Psychology, Flinders University, Adelaide, SA, Australia

^d^ Flinders Centre for Innovation in Cancer, Adelaide, SA, Australia

^e^ Centre for Neuroscience, Adelaide, SA, Australia

^f^ School of Biomedical Sciences and Pharmacy, University of Newcastle, Newcastle, NSW, Australia

^g^ School of Medical Sciences, University of New South Wales (UNSW Sydney), Sydney, NSW, Australia

Table of Contents

Supplementary Table 1. Correlated CpG sites (r > 0.1) within 500bp of each other.	2

Supplementary Table 2. Odds ratios and 95% Confidence Intervals (CI) for differentially methylated CpG sites among 1051 sites tested for association with schizophrenia (see Excel Supplementary Table 3 for the full results).	3

### Supplementary Table 1. Correlated CpG sites (r > 0.1) within 500bp of each other. 2

| Excluded from analysis | Retained in analysis |
| --- | --- |
| cg21113478 | cg00252934 |
| cg01956717 | cg22062554 |
| cg02387701 | cg10301588 |
| cg03680898 | cg03451959 |
| cg04232615 | cg04784327 |
| cg10798745 | cg12832565 |
| cg11003133 | cg27541604 |
| cg11236746 | cg20730966 |
| cg11271526 | cg13855862 |
| cg26509254 | cg16253551 |

### Supplementary Table 2. Odds ratios and 95% Confidence Intervals (CI) for differentially methylated CpG sites among 1051 sites tested for association with schizophrenia (see Excel Supplementary Table 3 for the full results).

| Probe ID | *B* | *OR* | 95% CI | *p* | FDR adj. *p* |
| --- | --- | --- | --- | --- | --- |
| cg06922393 | 0.019 | 1.019 | 1.008-1.030 | 0.001 | 0.331 |
| cg03035167 | 0.017 | 1.017 | 1.006-1.026 | 0.001 | 0.331 |
| cg12632097 | -0.007 | 0.993 | 0.988-0.997 | 0.001 | 0.331 |
| cg24897255 | -0.011 | 0.989 | 0.982-0.995 | 0.002 | 0.331 |
| cg09306340 | -0.009 | 0.991 | 0.986-0.996 | 0.002 | 0.331 |
| cg10833838 | 0.017 | 1.017 | 1.006-1.028 | 0.002 | 0.331 |
| cg11993160 | 0.020 | 1.020 | 1.006-1.033 | 0.003 | 0.331 |
| cg08446539 | 0.014 | 1.014 | 1.004-1.023 | 0.003 | 0.331 |
| cg10066189 | -0.005 | 0.995 | 0.991-0.998 | 0.003 | 0.331 |
| cg09682727 | 0.015 | 1.015 | 1.005-1.025 | 0.004 | 0.331 |
| cg09862509 | 0.011 | 1.011 | 1.003-1.019 | 0.004 | 0.331 |
| cg18143792 | 0.026 | 1.026 | 1.008-1.043 | 0.004 | 0.331 |
| cg07128021 | 0.015 | 1.015 | 1.004-1.024 | 0.005 | 0.375 |
| cg00089453 | 0.011 | 1.011 | 1.003-1.018 | 0.007 | 0.504 |
| cg21508639 | 0.012 | 1.012 | 1.003-1.021 | 0.009 | 0.582 |
| cg24862861 | -0.004 | 0.996 | 0.992-0.998 | 0.009 | 0.582 |
| cg13803727 | 0.012 | 1.012 | 1.003-1.021 | 0.009 | 0.582 |
| cg02979010 | -0.016 | 0.984 | 0.972-0.996 | 0.010 | 0.597 |
| cg00904966 | -0.026 | 0.974 | 0.954-0.993 | 0.011 | 0.597 |
| cg23292259 | -0.008 | 0.992 | 0.985-0.998 | 0.011 | 0.597 |
| cg19187564 | -0.011 | 0.989 | 0.980-0.997 | 0.014 | 0.614 |
| cg17813874 | 0.015 | 1.015 | 1.002-1.026 | 0.015 | 0.614 |
| cg20077042 | 0.009 | 1.009 | 1.001-1.015 | 0.015 | 0.614 |
| cg09146903 | -0.008 | 0.992 | 0.985-0.998 | 0.016 | 0.614 |
| cg13855862 | 0.009 | 1.009 | 1.001-1.016 | 0.016 | 0.614 |
| cg07430085 | 0.006 | 1.006 | 1.001-1.011 | 0.016 | 0.614 |
| cg05257202 | 0.016 | 1.016 | 1.002-1.029 | 0.017 | 0.614 |
| cg08966978 | -0.004 | 0.996 | 0.992-0.999 | 0.017 | 0.614 |
| cg24152264 | 0.013 | 1.013 | 1.002-1.023 | 0.018 | 0.614 |
| cg00676085 | 0.018 | 1.018 | 1.003-1.032 | 0.018 | 0.614 |
| cg12894371 | 0.019 | 1.020 | 1.003-1.036 | 0.019 | 0.614 |
| cg19411699 | 0.010 | 1.010 | 1.001-1.018 | 0.019 | 0.614 |
| cg14287546 | -0.004 | 0.996 | 0.992-0.999 | 0.020 | 0.614 |
| cg07816095 | 0.006 | 1.006 | 1.000-1.010 | 0.021 | 0.614 |
| cg10923036 | 0.010 | 1.010 | 1.001-1.018 | 0.023 | 0.614 |
| cg15508749 | -0.007 | 0.993 | 0.986-0.998 | 0.023 | 0.614 |
| cg05334655 | 0.010 | 1.010 | 1.001-1.019 | 0.024 | 0.614 |
| cg11860203 | 0.011 | 1.011 | 1.001-1.021 | 0.024 | 0.614 |
| cg01904296 | 0.009 | 1.009 | 1.001-1.017 | 0.025 | 0.614 |
| cg24469412 | -0.007 | 0.993 | 0.986-0.999 | 0.025 | 0.614 |
| cg25733915 | -0.004 | 0.996 | 0.992-0.999 | 0.025 | 0.614 |
| cg12551813 | 0.014 | 1.014 | 1.001-1.026 | 0.025 | 0.614 |
| cg21935292 | -0.004 | 0.996 | 0.991-0.999 | 0.027 | 0.614 |
| cg14592365 | 0.013 | 1.013 | 1.001-1.025 | 0.027 | 0.614 |
| cg08715914 | -0.005 | 0.995 | 0.990-0.999 | 0.027 | 0.614 |
| cg05995496 | 0.009 | 1.009 | 1.001-1.017 | 0.028 | 0.614 |
| cg25436157 | 0.012 | 1.012 | 1.001-1.022 | 0.029 | 0.614 |
| cg17237603 | 0.007 | 1.007 | 1.000-1.014 | 0.029 | 0.614 |
| cg22457668 | -0.010 | 0.990 | 0.980-0.998 | 0.029 | 0.614 |
| ch.5.95626522F | -0.002 | 0.998 | 0.995-0.999 | 0.031 | 0.614 |
| cg02586507 | 0.007 | 1.007 | 1.000-1.013 | 0.031 | 0.614 |
| cg01556246 | 0.013 | 1.013 | 1.001-1.024 | 0.033 | 0.614 |
| cg04656070 | -0.028 | 0.972 | 0.947-0.997 | 0.033 | 0.614 |
| cg22920792 | 0.004 | 1.004 | 1.000-1.008 | 0.033 | 0.614 |
| cg01717524 | 0.008 | 1.008 | 1.000-1.015 | 0.034 | 0.614 |
| cg17594003 | -0.033 | 0.968 | 0.938-0.997 | 0.034 | 0.614 |
| cg17824939 | -0.023 | 0.977 | 0.956-0.998 | 0.035 | 0.614 |
| cg24180759 | 0.011 | 1.011 | 1.000-1.021 | 0.035 | 0.614 |
| cg07627445 | 0.007 | 1.007 | 1.000-1.012 | 0.035 | 0.614 |
| cg11046602 | 0.014 | 1.015 | 1.001-1.028 | 0.035 | 0.614 |
| cg15032960 | 0.017 | 1.017 | 1.001-1.032 | 0.037 | 0.614 |
| cg03778707 | 0.006 | 1.006 | 1.000-1.012 | 0.037 | 0.614 |
| cg01199721 | -0.005 | 0.995 | 0.990-0.999 | 0.037 | 0.614 |
| cg07442040 | -0.010 | 0.990 | 0.980-0.999 | 0.038 | 0.614 |
| cg22117666 | 0.010 | 1.010 | 1.000-1.020 | 0.039 | 0.614 |
| cg23653444 | 0.005 | 1.005 | 1.000-1.009 | 0.039 | 0.614 |
| cg03923676 | -0.018 | 0.982 | 0.965-0.999 | 0.040 | 0.622 |
| cg14571714 | 0.004 | 1.004 | 1.000-1.008 | 0.041 | 0.630 |
| cg25462842 | 0.009 | 1.009 | 1.000-1.018 | 0.041 | 0.630 |
| cg08798307 | 0.011 | 1.011 | 1.000-1.021 | 0.043 | 0.639 |
| cg19548313 | 0.008 | 1.008 | 1.000-1.016 | 0.044 | 0.639 |
| cg22726026 | 0.003 | 1.003 | 1.000-1.006 | 0.044 | 0.639 |
| cg11594131 | -0.004 | 0.996 | 0.992-0.999 | 0.045 | 0.639 |
| cg17372806 | 0.015 | 1.015 | 1.000-1.030 | 0.045 | 0.639 |
| cg23200218 | 0.016 | 1.016 | 1.000-1.031 | 0.046 | 0.645 |
| cg07267600 | 0.009 | 1.009 | 1.000-1.018 | 0.048 | 0.655 |
| cg26385283 | 0.012 | 1.012 | 1.000-1.024 | 0.049 | 0.655 |
| cg06669036 | 0.004 | 1.004 | 1.000-1.008 | 0.049 | 0.655 |
| cg27541604 | 0.012 | 1.012 | 1.000-1.023 | 0.050 | 0.661 |
